# INGOT-DR: an interpretable classifier for predicting drug resistance in M. tuberculosis

**DOI:** 10.1101/2020.05.31.115741

**Authors:** Hooman Zabeti, Nick Dexter, Amir Hosein Safari, Nafiseh Sedaghat, Maxwell Libbrecht, Leonid Chindelevitch

## Abstract

**Motivation:** Prediction of drug resistance and identification of its mechanisms in bacteria such as *Mycobacterium tuberculosis*, the etiological agent of tuberculosis, is a challenging problem. Solving this problem requires a transparent, accurate, and flexible predictive model. The methods currently used for this purpose rarely satisfy all of these criteria. On the one hand, approaches based on testing strains against a catalogue of previously identified mutations often yield poor predictive performance; on the other hand, machine learning techniques typically have higher predictive accuracy, but often lack interpretability and may learn patterns that produce accurate predictions for the wrong reasons. Current interpretable methods may either exhibit a lower accuracy or lack the flexibility needed to generalize them to previously unseen data.

**Contribution:** In this paper we propose a novel technique, inspired by the group testing and Boolean compressed sensing, which yields highly accurate predictions, interpretable results, and is flexible enough to be optimized for various evaluation metrics at the same time.

**Results:** We test the predictive accuracy of our approach on five first-line and seven second-line antibiotics used for treating tuberculosis. We find that it has a higher or comparable accuracy to that of commonly used machine learning models, and is able to identify variants in genes with previously reported association to drug resistance. Our method is intrinsically interpretable, and can be customized for different evaluation metrics. Our implementation is available at github.com/hoomanzabeti/INGOT_DR and can be installed via The Python Package Index (Pypi) under **ingotdr**. This package is also compatible with most of the tools in the Scikit-learn machine learning library.

## Introduction

Drug resistance is the phenomenon by which an infectious organism (also known as pathogen) develops resistance to one or more drugs that are commonly used in treatment [1]. In this paper we focus our attention on *Mycobacterium tuberculosis* (MTB), the etiological agent of tuberculosis, which is the largest single infectious agent killer in the world today, responsible for over 10 million detected cases and approximately 1.4 million deaths only in 2019 [2].

The development of resistance to common drugs used in treatment is a serious public health threat, not only in low and middle-income countries, but also in high-income countries where it is particularly problematic in hospital settings [3]. It is estimated that, without the urgent development of novel antimicrobial drugs, the total mortality due to drug resistance will exceed 10 million people a year by 2050, a number exceeding the annual mortality due to cancer today [4].

Existing models for predicting drug resistance from whole-genome sequencing (WGS) data broadly fall into two classes. The first, which we refer to as “catalogue methods,” involves testing the WGS data of an isolate for the presence of point mutations (most often single-nucleotide polymorphisms, or SNPs) associated with known drug resistance. These mutations are typically identified via a microbial genome-wide association study (GWAS) and may be confirmed with a functional genomics study. If at least one previously identified mutation is present, the isolate is declared to be resistant [5, 6, 7, 8, 9]. While these methods are simple to understand and apply, they often suffer from poor predictive accuracy [10], especially in identifying new resistance mechanisms or predicting resistance to rarely used drugs.

The second class, which we will refer to as “machine learning methods”, seeks to infer the drug resistance of an isolate by training complex models directly on WGS and drug susceptibility test (DST) data [11, 12, 13]. Such methods tend to result in highly accurate predictions at the cost of flexibility and interpretability - specifically, they typically provide only limited, if any, insights into the drug resistance mechanisms involved, and often do not impose explicit limits on the predictive model’s complexity. Learning approaches based on deep neural networks [13, 14] are an example of very accurate but very complex “black-box” models of drug resistance.

In this paper we propose a novel method, based on the group testing problem [15] and Boolean compressed sensing (CS), for the prediction of drug resistance. CS is a mathematical technique for sparse signal recovery from under-determined systems of linear equations [16], and has been successfully applied in many application areas including digital signal processing [17, 18], MRI imaging [19], radar detection [20], and computational uncertainty quantification [21, 22]. Under a sparsity assumption on the unknown signal vector, it has been shown that CS techniques enable recovery from far fewer measurements than required by the Nyquist-Shannon sampling theorem [23]. Boolean CS is a slight modification of the CS problem, replacing linear algebra over the real numbers with a Boolean algebra over binary numbers [24], which has been successfully applied to various forms of non-adaptive group testing [24, 25, 26].

Our approach, INterpretable GrOup Testing for Drug Resistance (INGOT-DR), combines the flexibility and interpretability of catalogue methods with the accuracy of machine learning methods. More specifically, INGOT-DR is capable of recovering interpretable rules for predicting drug resistance that both result in a high classification accuracy as well as provide insights into the mechanisms of drug resistance. We compare the performance of INGOT-DR with that of standard and state-of-the-art machine learning and rule-based learning methods which have been previously used for genotype-phenotype prediction on MTB data. These methods are logistic regression (LR) [27], random forests (RF) [28], Support Vector Machines (SVM) [29], and KOVER [30]. The comparison covers the prediction of drug resistance for twelve drugs, of which five are first-line and seven are second-line drugs. INGOT-DR displays a competitive performance while maintaining interpretability, flexibility, and accurately recovering many of the known mechanisms of drug resistance.

## 1 Methods

We present our methodology as follows. Sections 1.1 and 1.2 introduce the group testing problem, and discuss how group testing can be combined with compressed sensing to deliver an interpretable predictive model. Section 1.3 introduces substantial modifications to a previously published method, which are needed to produce an accurate and flexible classifier that can be tuned for specific evaluation metrics and tasks. Section 1.4 describes the tuning process required to provide the desired trade-off between sensitivity and specificity in a model’s predictions.

### 1.1 Group testing and Boolean compressed sensing

We frame the problem of predicting drug resistance given sequence data as a group testing problem, originally introduced in [15]. This approach for detecting defective members of a set was motivated by the need to screen a large population of soldier recruits for syphilis in the United States during the World War II. The screening, performed by testing blood samples, was costly due to the low numbers of infected individuals. To make the screening more efficient, Robert Dorfman suggested pooling blood samples into specific groups and testing the groups instead. A positive result for the group would imply the presence of at least one infected member. The problem then becomes one of finding the subset of individuals whose infected status can explain all the positive results without invalidating any of the negative ones.

In this setting, the **design matrix** encodes the individuals tested in each group, the **outcome vector** describes the result of each test, and the solution, obtained from a suitable algorithmic procedure, is a {0, 1}-valued vector representing the infection status of the individuals [24, 31]. Since the fraction of infected individuals is assumed to be small, the solution vector is sparse and can be recovered using relatively few tests with Boolean CS. The importance of this observation lies in the fact that the result of solving the Boolean CS problem can also be interpreted as a sparse set of rules for determining the status of each sample in other data mining contexts [24]. We summarize this correspondence in our context in Table 1 below, and use the context-specific interpretation throughout the rest of this paper.

**Table 1.**
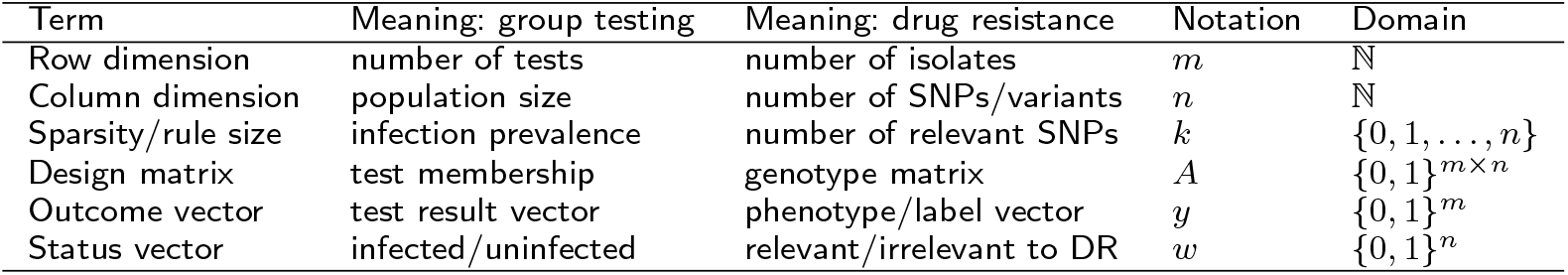
Correspondence between group testing and the drug resistance prediction problem.

Mathematically, the problem with *m* isolates and *n* SNPs can be described by the Boolean design matrix *A* ∈ {0, 1}^*m*×*n*^, where *A*_*ij*_ indicates the presence/absence status of SNP *j* in the *i*-th isolate, and the Boolean outcome vector *y* ∈ {0, 1}^*m*^, where *y*_*i*_ represents the drug resistance phenotype of the *i*-th isolate. Let us define the relevance vector *w* ∈ {0, 1}^*n*^ in such a way that *w*_*j*_ = 1 if and only if the *j*-th SNP is relevant to drug resistance.

The key assumption is that one or more SNPs relevant to drug resistance can cause the isolate to be drug-resistant, whereas an isolate with no such SNP will be drug-sensitive. This is an assumption commonly made in the literature, and is precisely the same as the key assumption of group testing, which is that the presence of one or more infected individuals leads to a positive test, while a test with no infected individuals comes out negative (we note that these assumptions only hold in the absence of noise). In fact, although our group is the first one, to our knowledge, to make the connection between group testing and drug resistance prediction, a previously published method for this task [32] corresponds almost perfectly to the Definite Defectives algorithm used in group testing [33].

Under this assumption, the outcome vector satisfies the relationship

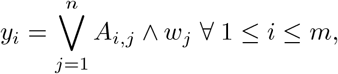

where ∨ and ∧ are the Boolean OR and AND operators, respectively. Using the definition of Boolean matrix-vector multiplication, this can be equivalently written

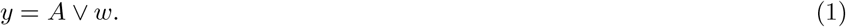

If the status vector *w* satisfying equation (1) is assumed to be sparse (i.e. there are few relevant SNPs), the problem of finding *w* becomes an instance of the sparse Boolean vector recovery problem:

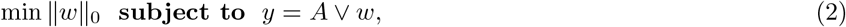

where ‖*w*‖ _0_, called the *𝓁*_0_ -norm of *w*, is the number of non-zero entries it contains.

The combinatorial optimization problem (2) is well-known to be NP-hard; see, e.g., [16, Section 2.3]. In [24, 34] an equivalent formulation of (2) via 0-1 integer linear programming (ILP) is proposed, in which the *𝓁*_0_-norm is replaced by the convex *𝓁*_1_-norm, equivalent to it over binary vectors, and the Boolean matrix-vector product is replaced with equivalent linear constraints. We recapitulate their formulation here:

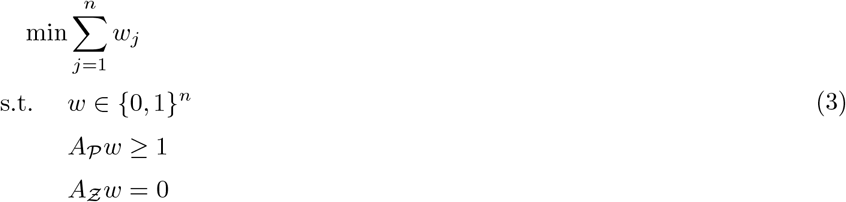

Here, 𝒫 := {*i* : *y*_*i*_ = 1} and *Ƶ* := {*i* : *y*_*i*_ = 0} are the sets of positive (drug-resistant) and negative (drug-sensitive) isolates, respectively, and *A*_*S*_ denotes the submatrix of *A* whose row indices are in the given subset *S*. In this formulation, the objective is to minimize the number of SNPs inferred to be relevant to drug resistance. The first constraint then ensures that each SNP is classified as either relevant or irrelevant, the second one ensures that the drug-resistant isolates have at least one relevant SNP present, and the third one ensures that the drug-sensitive isolates do not have any such SNPs, in line with our key assumption. This NP-hard problem formulation can further be made tractable for linear programming by relaxing the Boolean constraint on *w* in (3) to 0 ≤ *w*_*j*_ ≤ 1 for all 1 ≤ *j* ≤ *n* [24].

Because the Boolean CS problem is based on Boolean algebra, the conditions on the Boolean matrices *A* that guarantee exact recovery of *k*-sparse status vectors (vectors with at most *k* 1’s) via such linear programming relaxations are quite stringent, and differ from those of standard CS. Specifically, in order to hold, these guarantees require the matrix *A* to be *k*-disjunct, i.e. for any Boolean sum of at most *k* of its columns to not be greater than or equal to any other column. As we have no control over *A* in our setting, no such recovery guarantees can be provided.

In [24], the combinatorial problem (3) is augmented with slack variables and a regularization term to trade off between the sparsity of *w* on the one hand, and the discrepancy between the predicted and the actual outcome vector on the other hand. With these modifications, the formulation becomes:

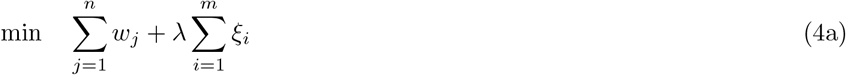

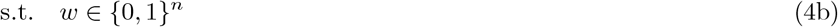

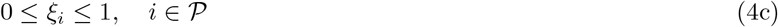

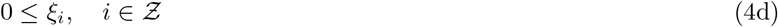

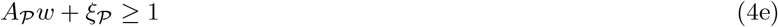

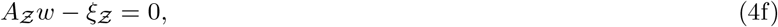

where *λ >* 0 is a regularization parameter and *ξ* is the so-called slack vector. Taking this formulation as a starting point, we introduce several refinements in Section 1.3.

### 1.2 From group testing to interpretable classification

As described in the previous section, the solution to the ILP (4) can be seen as an interpretable rule-based classifier in contexts beyond group testing. The status vector *w* naturally encodes the following rule: If any feature *f* with *w*_*f*_ = 1 is present in the sample, classify it as positive; otherwise, classify it as negative. More formally, assume that we have a labelled dataset 𝒟 = {(*x*_1_, *y*_1_), …, (*x*_*m*_, *y*_*m*_)}, where the *x*_*i*_ ∈ 𝒳 := {0, 1}^*n*^ are *n*-dimensional binary feature vectors and the *y*_*i*_ ∈ {0, 1} are the binary labels. The feature matrix *A* is defined via *A*_*i,j*_ = (*x*_*i*_)_*j*_ (the *j*-th component of the *i*-th feature vector). If *ŵ* is the solution of ILP (4) for this matrix *A* and the outcome vector 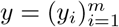, we define the classifier *ĉ* : 𝒳 → {0, 1} via

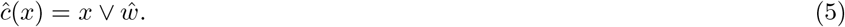

What makes this classifier interpretable is that it explicitly depends on the presence or absence of specific features in its input, while ignoring all the other features.

### 1.3 Our approach leads to a refined ILP formulation

The formulation of the ILP (4) is designed to provide, via the parameter *λ*, a trade-off between the sparsity of a rule and the total slack, a quantity that resembles (but does not equal) the training error. We now describe a refinement of this formulation that directly encodes the different types of error, which provides more flexibility during the training process by allowing us to optimize a more precise objective function that is particularly suitable to the application at hand.

As was done in the previous section, we assume that *ĉ* is the binary classifier obtained by training with a Boolean feature matrix *A* and its corresponding label vector *y*. We further refer to a misclassified training sample as a **false negative** if it has label 1 (is in 𝒫), and as a **false positive** if it has label 0 (is in *Ƶ*). In the drug resistance setting, a false negative would mean that we incorrectly predict a drug-resistant isolate to be drug-sensitive, while a false positive would mean that we predict a drug-sensitive isolate to be drug-resistant.

We begin by noting that in the ILP (4), each entry of *ξ*_𝒫_ must take on a value of 0 or 1, and a value of 1 corresponds to a false negative for *ĉ*. This follows from the fact that *A* is a binary matrix and *w* is a binary vector, so the optimal *ξ*_𝒫_ is also a binary vector (since *λ >* 0), and therefore

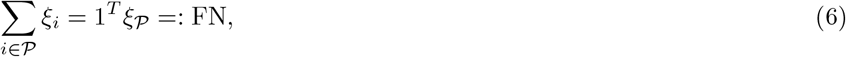

where we use FN to denote the number of false negatives.

However, *ξ*_*Ƶ*_ in the ILP (4) can take on integer values greater than 1 corresponding to false positives for *ĉ*. To be able to express the number of false positives, denoted FP, we modify the constraints (4d) and (4f) by also setting

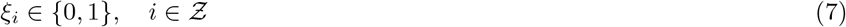

and replacing the equality constraint *A*_*Ƶ*_ *w* −*ξ*_*Ƶ*_ = 0 with the tighter inequality

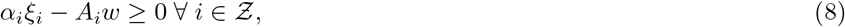

where 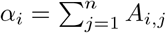 and *A*_*i*_ is the *i*th row of *A*.

After these modifications, (8) ensures that *ξ*_*i*_ = 1 if *A*_*i*_*w >* 0, while the presence of *ξ*_*Ƶ*_ in the objective function, with *λ >* 0, ensures that *ξ*_*i*_ = 0 if *A*_*i*_*w* = 0, for any *i* ∈ *Ƶ*. We now also get

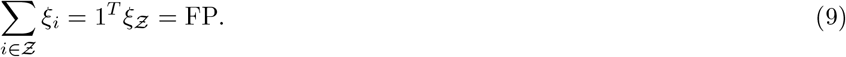

To provide additional flexibility for situations where false positives and false negatives are valued differently, we further split the regularization term into two: one for the positive class 𝒫, and one for the negative class *Ƶ*:

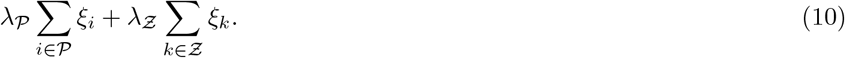

The general form of the new ILP is now as follows:

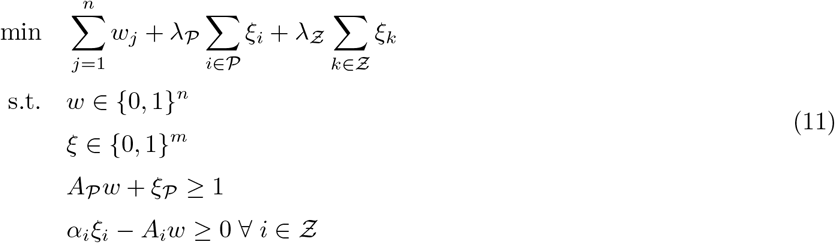

In this new formulation, *λ*_𝒫_ and *λ*_*Ƶ*_ control the trade-off between the false positives and the false negatives, and jointly influence the sparsity of the rule. In the following section we describe how this formulation can be further tailored to optimize different evaluation metrics, such as the sensitivity and the specificity of the predictor.

### 1.4 Optimizing different target metrics such as the sensitivity and the specificity

Since the ILP formulation in (11) provides us with direct access to the two components of the training error as well as the sparsity (rule size), we may modify the classifier to optimize a variety of target metrics by transforming some of the objective function components into constraints and optimizing the remaining ones.

For instance, assume that we would like to train the classifier *ĉ* to maximize the sensitivity at a given minimum specificity 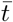 and maximum rule size *k*. Recall that

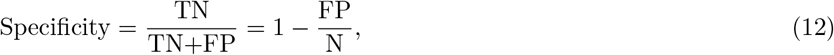

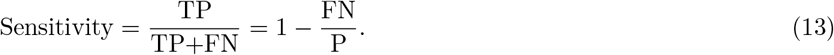

From equation (10), equation (12) and the definition of *Ƶ*, we get the constraint

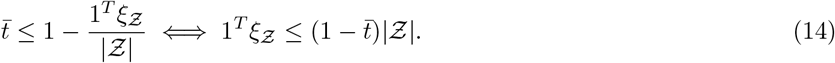

Also, to restrict the maximum rule size to *k* we can use the constraint

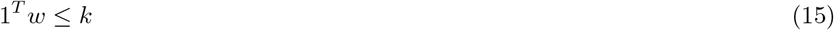

Our objective is to maximize the sensitivity, which is equivalent to minimizing

∑_*i*∈𝒫_ *ξ*_*i*_ by equations (13) and (6). In addition, by adding equations (14) and (15), the ILP (11) can be modified as follows:

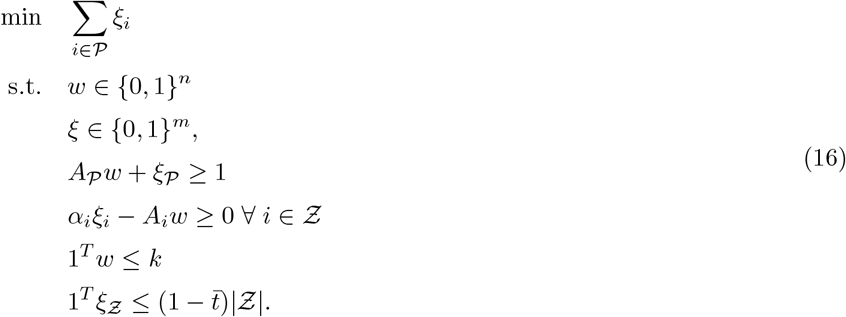

The maximum specificity at given sensitivity and rule size can be found analogously. In a similar way, one can minimize a weighted average of rule size and false positive rate at a given maximum false negative rate (minimum sensitivity), or vice versa.

## 2 Implementation

### 2.1 Existing methods used for comparison with INGOT-DR

To ensure a fair comparison, we use three popular machine learning methods used for drug resistance prediction: random forests (RF) [28], logistic regression (LR) [27], and support vector machines (SVM) [29]. The use of RF is motivated by its flexibility and its many successful applications in computational biology and genomics [35, 36]. The use of LR is based on its excellent performance in drug resistance prediction for MTB in comparison to other methods [37]. The use of SVM is motivated by its excellent performance in a comparison of drug resistance prediction for multiple bacterial pathogens [30]; we use it with a linear kernel for simplicity, although other kernels are often used [38]. For LR and SVM, we consider the *𝓁*_1_ and *𝓁*_2_ regularizations, which correspond to penalizing the sum of the absolute values and the Euclidean norm of the coefficients, respectively.

We also use, to our knowledge, the only other interpretable machine learning method for drug resistance prediction, KOVER [30]. All the methods except KOVER are implemented in the Python programming language [39]. Although KOVER can provide rule-based classifiers from two algorithms: Classification and Regression Trees (CART) and Set Covering Machine (SCM), we only consider the latter as it is the main innovation of KOVER [40], and the two algorithms yield very similar accuracy [30]. We use the Scikit-learn [41] implementation for the machine learning models - *RandomForestClassifier* for RF, *LogisticRegression* for LR, and *LinearSVC* for SVM. We also use KOVER version 2.0 [42], and harness the Python API to the CPLEX optimizer, version 12.10.0 [43], through the Pulp API [44, 45] to solve the ILPs in INGOT-DR.

### 2.2 Data

We combine data from the Pathosystems Resource Integration Center (PATRIC) [46] and the Relational Sequencing TB Data Platform (ReSeqTB) [47]. This results in 8,000 isolates together with their resistant/susceptible status for twelve drugs, including five first-line (rifampicin, isoniazid, pyrazinamide, ethambutol, and streptomycin) and seven second-line drugs (kanamycin, amikacin, capreomycin, ofloxacin, moxifloxacin, ciprofloxacin, and ethionamide) [48, 49]. The whole-genome sequencing data for these 8,000 isolates, in the form of paired FASTQ files, are downloaded from the European Nucleotide Archive [50] and the Sequence Read Archive [51]. The accession numbers used to obtain the data in our study are: ERP[000192, 000520, 006989, 008667, 010209, 013054], PRJEB[10385, 10950, 14199, 2358, 2794, 5162, 9680], PRJNA[183624, 235615, 296471], and SRP[018402, 051584, 061066].

In order to transform the raw sequencing data into variant calls, we use a pipeline similar to that used in previous work [52, 49]. We use the BWA software [53], specifically, the BWA-MEM program, for the mapping. We then call the single-nucleotide polymorphisms (SNPs) of each isolate with two different pipelines, SAMtools [54] and GATK [55], and take the intersection of their calls to ensure reliability. The final dataset, which includes the position as well as the reference and alternative allele for each SNP [49], is used as the input to our machine learning tools.

Starting from this input we create a binary feature matrix as described in Section 1.2. For each drug, we only consider the isolates with a status for this drug. We group all the SNPs in perfect linkage disequilibrium (LD) [56], i.e. sharing identical presence/absence patterns in those isolates, into a single feature that we call a SNP group. This representation does not affect the predictive accuracy of any machine learning methods, but helps create a consistent feature importance score for the non-interpretable ones. In KOVER, at most one SNP in a SNP group can be selected to be part of a rule, and the remaining SNPs in the group are labelled equivalent [40]; we adopt this convention here. The number of labeled and drug-resistant isolates, as well as the number of SNPs and SNP groups for each drug, is shown in Table 2.

**Table 2.** Summary statistics for our dataset, with a line separating first-line and second-line drugs.

| Drug | # of isolates | # of resistant isolates | # of SNPs | # of SNP groups |
| --- | --- | --- | --- | --- |
| Ethambutol | 6,096 | 1,407 | 597,133 | 55,164 |
| Isoniazid | 7,734 | 3,445 | 642,373 | 65,090 |
| Pyrazinamide | 3,858 | 754 | 281,432 | 33,942 |
| Rifampicin | 7,715 | 2,968 | 646,855 | 65,379 |
| Streptomycin | 5,125 | 2,104 | 542,640 | 45,037 |
| Kanamycin | 2,436 | 697 | 391,708 | 21,513 |
| Amikacin | 2,033 | 573 | 141,952 | 17,103 |
| Capreomycin | 1,991 | 552 | 341,935 | 15,389 |
| Ofloxacin | 2,911 | 800 | 407,235 | 23,905 |
| Moxifloxacin | 961 | 129 | 97,700 | 11,927 |
| Ciprofloxacin | 443 | 37 | 43,950 | 5,563 |
| Ethionamide | 1,516 | 498 | 344,960 | 15,145 |

### 2.3 Splitting the data into a training and testing set; tuning the hyper-parameters

To evaluate our classifier we use a random stratified train-test split, where the training set contains 80% and the testing set contains 20% of data. For hyper-parameter tuning, Scikit-learn provides two main approaches: grid search and randomized search cross-validation. KOVER is also equipped with two tuning techniques, *K*-fold cross-validation and risk bound selection. To make the comparison as consistent as possible, we use 5-fold cross-validation for KOVER and grid search with 5-fold cross-validation for all the other models. During cross-validation, balanced accuracy is used as the model selection metric for all the models except KOVER; to the best of our knowledge, KOVER does not provide the option to change the model selection metric.

### 2.4 Evaluating the models’ performance

Evaluating the performance of an interpretable predictive model can be challenging. While most evaluation methods focus on predictive accuracy, it is essential to assess the model’s interpretability as well. Although there is no consensus definition of interpretability, [57] suggest that an interpretable method should be able to provide an acceptable predictive accuracy while being easy to understand and provide meaningful insights to its audience. Adopting their idea, we evaluate the performance of our approach and the competitor methods using three metrics:

1. Predictive accuracy, measured via the balanced accuracy,

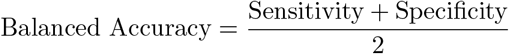
2. Simplicity, measured via the number of features (SNPs) in the trained model.
3. Insight generation, measured via the relevance of the selected SNPs to known drug resistance mechanisms.

This evaluation process is demonstrated in detail in Sections 2.5 and 3.

### 2.5 The comparison between interpretable and non-interpretable models

The overall pipeline consists of SNP calling and SNP grouping as described in Section 2.2, hyper-parameter tuning as described in Section 2.3, and model training and testing using the balanced accuracy as the metric as described in Section 2.4. This addresses the first evaluation criterion, the predictive accuracy.

To evaluate model simplicity, we investigate the SNPs selected by each model. For the rule-based classifiers, we ensure a low model complexity, and therefore a higher interpretability, by training both INGOT-DR and KOVER with the same maximum allowed rule size (number of SNPs used), *k*. By default, INGOT-DR also has a (training) specificity lower bound of 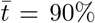, via the constraint explained in Section 1.4. We evaluate the simplicity of the remaining models by counting the SNPs with non-zero coefficients for LR and SVM, and the SNPs with a non-zero importance according to Scikit-learn for RF.

Lastly, to evaluate and fairly compare the models’ ability to generate insights, we compare the top *k* most important SNPs for each one [58]. For both INGOT-DR and KOVER, we simply evaluate the *k* or fewer SNPs used in each rule. Since the other machine learning methods are not inherently interpretable, we extract the SNP importance values using the Shapley additive explanation (SHAP) algorithm [59], a model-agnostic method for making explainable predictions rooted in game theory. This algorithm, implemented in the SHAP Python package, version 0.37.0 [60], provides the guaranteed unique solution satisfying three fairness conditions. We apply *TreeExplainer* for RF and *LinearExplainer* for LR and SVM, and select the *k* SNPs with the highest importance. We use *k* = 20 in all our experiments.

## 3 Results

### 3.1 INGOT-DR produces accurate predictive models

The performance of INGOT-DR compared to that of the other methods in terms of the balanced accuracy is summarized in Table 3, and Figure 1 separately shows the sensitivity and specificity. Overall, INGOT-DR outperforms all other models on 4/12 of the drugs, obtains the best performance (tied with KOVER) on an additional drug, and achieves a balanced accuracy within 5% of the best one for the remaining 7/12 drugs. SVM-l1 achieves the best balanced accuracy in 4/12 of the drugs, while LR-l1 and KOVER obtain the best balanced accuracy in 2/12 drugs each. Furthermore, INGOT-DR has a performance exceeding that of that of RF in 12/12 drugs, that of KOVER, LR-l2, and SVM-l2 in 9/12 drugs, that of LR-l1 in 8/12 drugs. SVM-l1 is the only competitive model, whose performance it only exceeds in 5/12 drugs, although it does obtain a marginally better balanced accuracy on average (85.7% vs 85.3%).

**Table 3.**
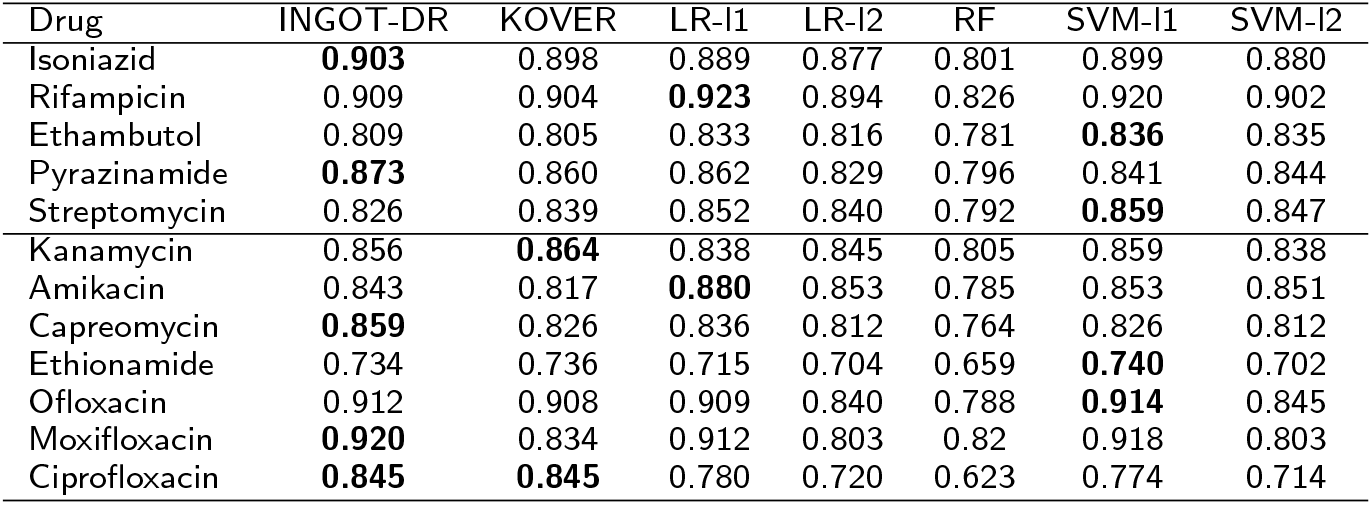
Balanced accuracy of all the methods in predicting drug resistance to 12 drugs.

| Drug | INGOT-DR | KOVER | LR-l1 | LR-l2 | RF | SVM-l1 | SVM-l2 |
| --- | --- | --- | --- | --- | --- | --- | --- |
| Isoniazid | **0.903** | 0.898 | 0.889 | 0.877 | 0.801 | 0.899 | 0.880 |
| Rifampicin | 0.909 | 0.904 | **0.923** | 0.894 | 0.826 | 0.920 | 0.902 |
| Ethambutol | 0.809 | 0.805 | 0.833 | 0.816 | 0.781 | **0.836** | 0.835 |
| Pyrazinamide | **0.873** | 0.860 | 0.862 | 0.829 | 0.796 | 0.841 | 0.844 |
| Streptomycin | 0.826 | 0.839 | 0.852 | 0.840 | 0.792 | **0.859** | 0.847 |
| Kanamycin | 0.856 | **0.864** | 0.838 | 0.845 | 0.805 | 0.859 | 0.838 |
| Amikacin | 0.843 | 0.817 | **0.880** | 0.853 | 0.785 | 0.853 | 0.851 |
| Capreomycin | **0.859** | 0.826 | 0.836 | 0.812 | 0.764 | 0.826 | 0.812 |
| Ethionamide | 0.734 | 0.736 | 0.715 | 0.704 | 0.659 | **0.740** | 0.702 |
| Ofloxacin | 0.912 | 0.908 | 0.909 | 0.840 | 0.788 | **0.914** | 0.845 |
| Moxifloxacin | **0.920** | 0.834 | 0.912 | 0.803 | 0.82 | 0.918 | 0.803 |
| Ciprofloxacin | **0.845** | **0.845** | 0.780 | 0.720 | 0.623 | 0.774 | 0.714 |

**Figure 1.**
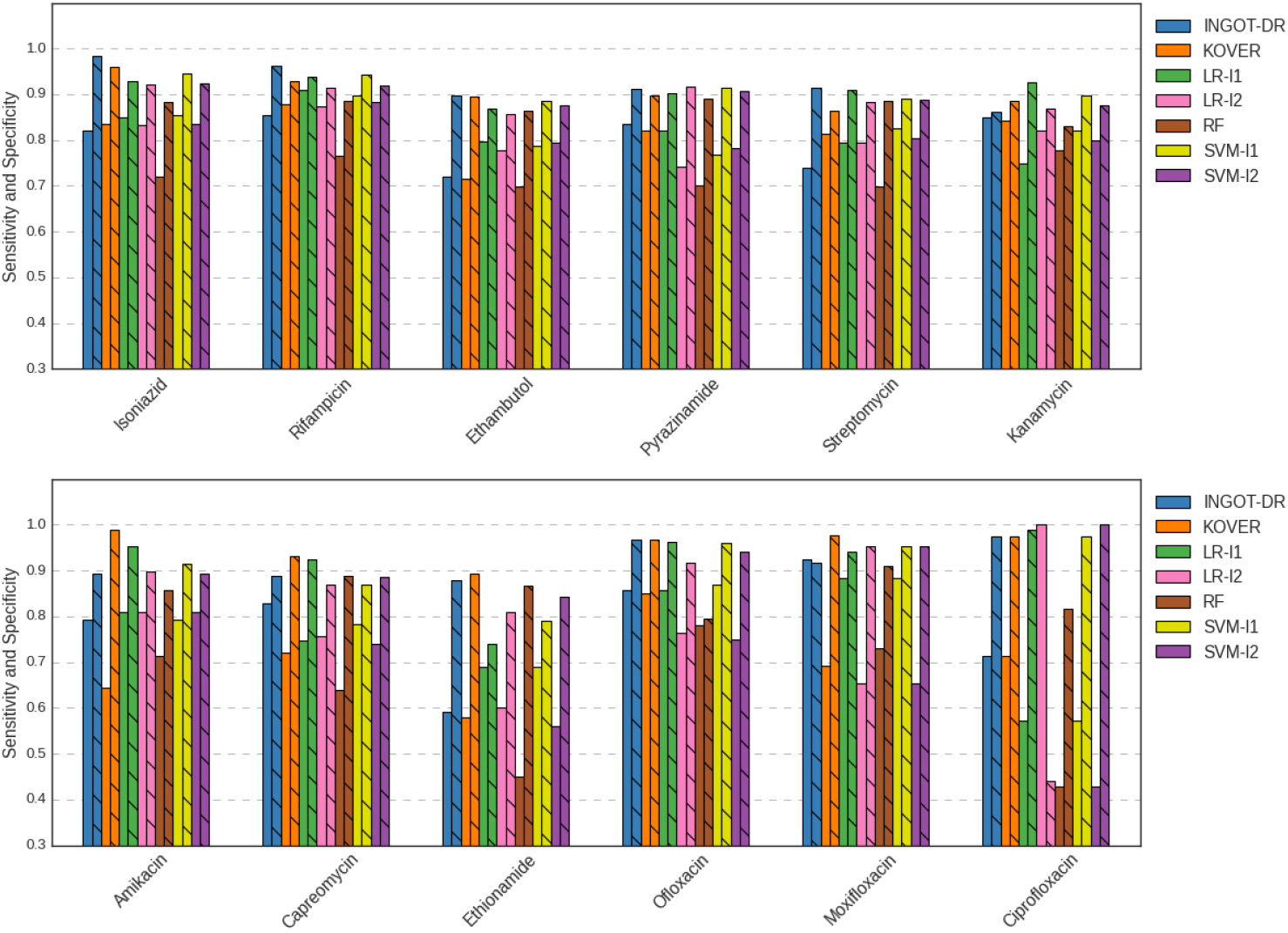
Sensitivity and specificity of all the methods in predicting drug resistance to 12 drugs.

### 3.2 INGOT-DR produces interpretable models

INGOT-DR produces predictive models in the form of disjunctive (logical-OR) rules over the presence of specific SNPs, as explained in Section 1.2. These models are easy to understand and interpret. Although KOVER considers rules containing both presence and absence of features [30], the absence of a SNP is harder to interpret in the context of genomics, so we only focus on the presence of SNPs here. We note that, by DeMorgan’s law, both methods could produce conjunctive (logical-AND) rules by training the model on the complement of the feature matrix, *Ā*, and outcome vector, 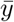; however, we focus on disjunctive rules in this paper.

We display the number of SNPs used by the predictive models produced by each method in Table 4. These results, combined with those of the previous section, suggest that INGOT-DR is producing the most interpretable models without sacrificing predictive accuracy. Although KOVER almost always produces shorter rules, they tend to not generalize as well to the testing dataset.

**Table 4.** Number of SNPs involved in the prediction made by each model for each drug.

| Drug | INGOT-DR | KOVER | LR-I1 | LR-I2 | RF | SVM-I1 | SVM-I2 |
| --- | --- | --- | --- | --- | --- | --- | --- |
| Isoniazid | 20 | 20 | 1,045 | 62,707 | 22,336 | 626 | 54,630 |
| Rifampicin | 20 | 20 | 739 | 63,621 | 29,373 | 476 | 52,732 |
| Ethambutol | 20 | 19 | 154 | 53,476 | 19,864 | 661 | 43,094 |
| Pyrazinamide | 20 | 17 | 114 | 32,885 | 9,495 | 428 | 25,485 |
| Streptomycin | 20 | 13 | 5,804 | 43,771 | 23,996 | 594 | 40,183 |
| Kanamycin | 20 | 20 | 2,383 | 20,934 | 9,314 | 231 | 18,716 |
| Amikacin | 20 | 19 | 2,252 | 16,622 | 7,639 | 212 | 14,260 |
| Capreomycin | 20 | 20 | 2,103 | 14,907 | 7,881 | 234 | 13,432 |
| Ethionamide | 20 | 20 | 41 | 14,791 | 7,777 | 280 | 13,551 |
| Ofloxacin | 20 | 17 | 394 | 23,206 | 14,312 | 265 | 19,694 |
| Moxifloxacin | 12 | 7 | 29 | 11,678 | 1,371 | 125 | 10,237 |
| Ciprofloxacin | 5 | 5 | 18 | 5,448 | 325 | 29 | 4,343 |

For a specific example, we consider the most concise model produced by INGOT-DR - the one for ciprofloxacin, a drug in the fluoroquinolone family. This model has a rule size of 5, and the SNPs used are all in the *gyrA* gene, known to be involved in the resistance to fluoroquinolones such as ciprofloxacin in bacteria [61]. In this example, INGOT-DR not only identifies the correct gene, but also selects mutations that are known to be associated with fluoroquinolone resistance in MTB - the selected codons, 90, 91 and 94, are among the codons most strongly associated with this type of resistance [62]. We state the rule obtained by INGOT-DR below, in a standard format specifying the gene, the original amino acid, the codon number, and the mutated amino acid.

**IF** gyrA A90V ∨ gyrA S91P ∨ gyrA D94A ∨ gyrA D94G ∨ gyrA D94Y **THEN** Resistant to ciprofloxacin

### 3.3 INGOT-DR selects many SNPs in genes previously associated with drug resistance

Our results demonstrate that the models produced by INGOT-DR contain many SNPs in genes previously associated with drug resistance in MTB. This suggests that INGOT-DR not only makes accurate predictions, but that it makes them for the right reason, and could thus also be used to prioritize hypotheses about the mechanisms associated with drug resistance.

Figure 2 shows, for each of the models, the top *k* ≤ 20 most important SNPs, defined as all the SNPs included in a rule by KOVER and INGOT-DR, and the top *k* SNPs by feature importance as defined by SHAP for the other models. We categorize each SNP according to the known information about its association with resistance to the drug of interest in MTB. This categorization is based on a list of 183 genes and 19 promoter regions selected out of over 4,000 MTB genes through a data-driven and consensus-driven process by a panel of experts [63]. We use the following categories:

**Figure 2.**
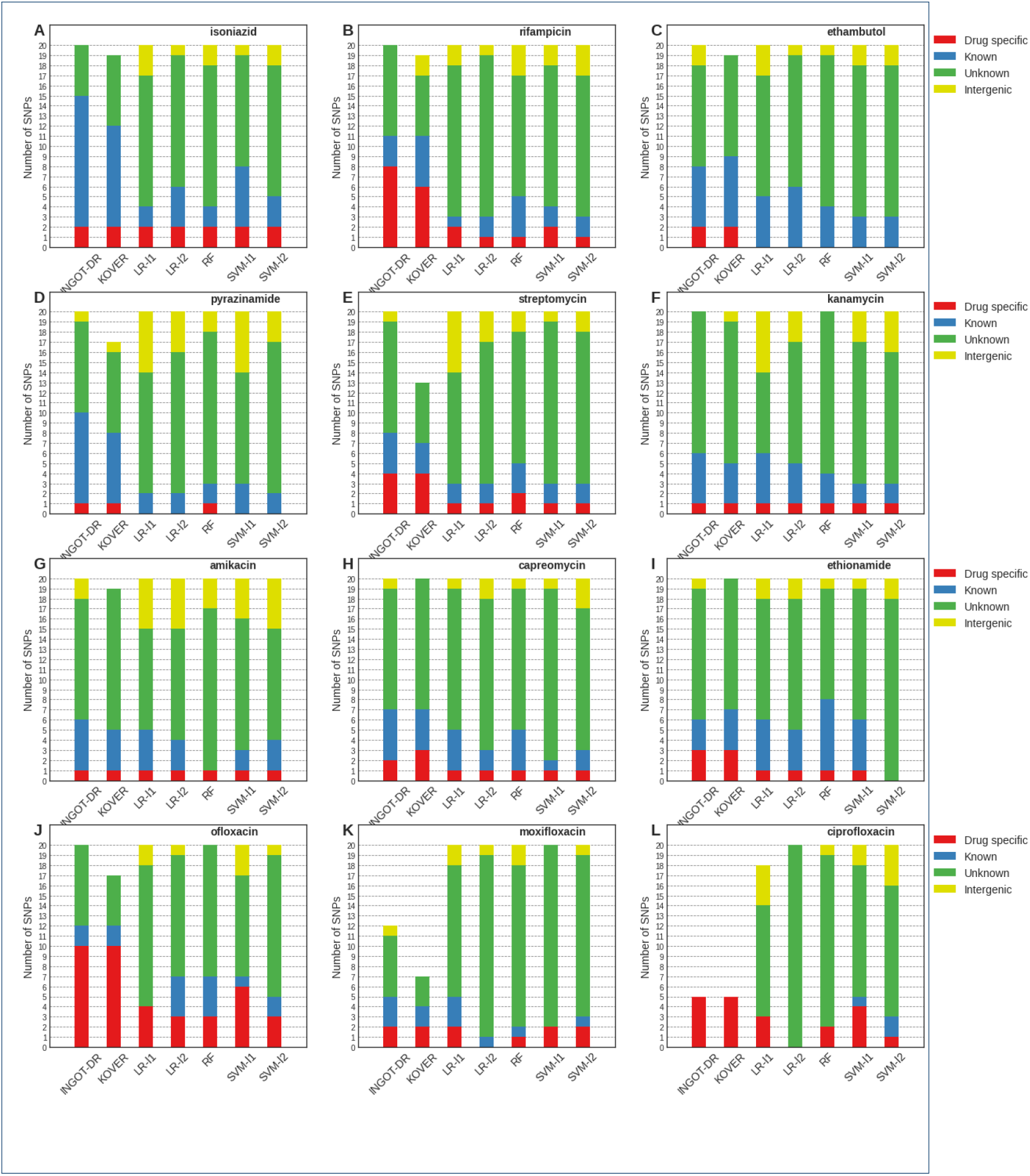
Top *k* ≤ 20 SNPs chosen by each model, categorized by association with drug resistance.

1. **Drug specific** association: SNP in a gene or intergenic region associated with drug resistance to the drug of interest;
2. **Known** association: SNP in a gene or intergenic region associated with drug resistance to any other drug;
3. **Unknown** association: SNP in a gene not known to be associated with drug resistance to any drug;
4. **Intergenic** association: SNP in an intergenic region not known to be associated with drug resistance to any drug.

We note that for the purposes of this categorization, whenever a group of SNPs in perfect LD was selected by the model, it was categorized according to the highest (lowest-numbered) category of any of the SNPs contained in the group. However, very few such SNP groups were selected by any of the models, and the absolute majority of the ones that were contained SNPs within the same gene.

A comparison between the methods based on Figure 2 suggests that INGOT-DR and KOVER detect more SNPs in regions known to be associated with drug resistance than all the other methods, with INGOT-DR detecting slightly more such SNPs than KOVER on average, even after adjusting for the slightly more concise rules produced by KOVER relative to INGOT-DR. However, with the exception of the most common first-line drugs (top row) and the three fluoroquinolones (bottom row), even the interpretable methods tend to select more SNPs in parts of the genome not known to be associated with drug resistance, suggesting the potentially important effects of population structure in MTB.

## 4 Conclusion

In this paper, we introduced a new approach for creating rule-based classifiers. Our method, INGOT-DR, utilizes techniques from group testing problem and Boolean compressed sensing, and leverages a 0 −1 ILP formulation. It produces classifiers that combine high accuracy with interpretability, and are flexible enough to be tailored for specific evaluation metrics.

We used INGOT-DR to produce classifiers for predicting drug resistance in MTB, by setting a minimum specificity of 90% and a maximum rule size of 20. We tested the classifiers’ predictive accuracy on a variety of antibiotics commonly used for treating tuberculosis, including five first-line and seven second-line drugs. We showed that INGOT-DR produces classifiers with a balanced accuracy exceeding that of other state-of-the-art rule-based and machine learning methods. In addition, we showed that INGOT-DR produces accurate models with a rule size small enough to keep the model understandable for human users. Finally, we showed that our approach generates insights by successfully identifying SNPs associated with drug resistance, as we ascertained on the specific example of ciprofloxacin.

We note that the presence of SNPs in perfect linkage disequilibrium (LD) [56], i.e. sharing identical presence/absence patterns, is common in bacteria such as MTB whose evolution is primarily clonal [64]. For this reason, while the grouping of such SNPs substantially simplifies the computational task at hand and makes it tractable, ascertaining the exact representative of each group to be selected to predict the drug resistance status of an isolate remains difficult. The presence of clonal structure within bacterial populations is a key challenge for the prediction of drug resistance, which we plan to address in future work.

In conclusion, our work has introduced a novel method, INGOT-DR, based on group testing techniques, for producing interpretable models of drug resistance, which demonstrated a state-of-the-art accuracy, descriptive ability, and relevance on an MTB dataset. In future work, we plan to address the challenges of population structure and to extend this framework to other bacteria as well as to less frequently used antimicrobial drugs. We expect our method to become a key part of the drug resistance prediction toolkit for clinical and public health microbiology researchers.

## Acknowledgements

The authors would like to acknowledge Dr. Cedric Chauve, Dr. Ben Adcock and Matthew Nguyen for helpful discussions, and the feedback from Dr. Nicholas Croucher, Dr. John Lees, Dr. Tim Walker, and Dr. Zamin Iqbal.

## Funding

This project was funded by a Genome Canada grant, “Machine learning to predict drug resistance in pathogenic bacteria”. LC acknowledges funding from the MRC Centre for Global Infectious Disease Analysis (MR/R015600/1), funded by the UK Medical Research Council (MRC) and the UK Foreign, Commonwealth & Development Office (FCDO) under the MRC/FCDO Concordat agreement, and is part of the EDCTP2 program supported by the EU.

### Abbreviations

CS: compressed sensing
FN: false negatives
FP: false positives
GWAS: genome-wide association study
ILP: integer linear programming
LD: linkage disequilibrium
LR: logistic regression
MTB: *Mycobacterium tuberculosis*
RF: random forests
SCM: set covering machine
SNP: single-nucleotide polymorphism
SVM: support vector machine
WGS: whole-genome sequencing

## Availability of data and materials

The data used in this study is freely available from the ENA and the NCBI. The code is freely available on GitHub.

## Ethics approval and consent to participate

No ethics approval was required for this study.

## Consent for publication

All authors have consented to this publication.

## Authors’ contributions

HZ has conceptualized and implemented the method, carried out the experiments, and analyzed the results. ND has contributed to the conceptualization and implementation. AS and NF have contributed to the data collection, preprocessing and analysis. ML and LC have contributed to conceptualizing the method, and supervised the research. All the authors have additionally contributed to writing or editing the draft and the final version of the manuscript.

